# Enhancing functionalized liposome avidity to cells via lipid phase separation

**DOI:** 10.1101/2022.10.18.512758

**Authors:** Timothy Q. Vu, Lucas E. Sant’Anna, Neha P. Kamat

**Author notes:** These authors contributed equally to this work (Timothy Q. Vu, Lucas E. Sant’Anna).

## Abstract

The addition of both cell-targeting moieties and polyethylene glycol (PEG) to nanoparticle (NP) drug delivery systems is a standard approach to improve the biodistribution, specificity, and uptake of therapeutic cargo. The spatial presentation of these molecules affects avidity of the NP to target cells in part through an interplay between the local ligand concentration and the steric hindrance imposed by PEG molecules. Here, we show that lipid phase separation in nanoparticles can modulate liposome avidity by changing the proximity of PEG and targeting protein molecules on a nanoparticle surface. Using lipid-anchored nickel-nitrilotriacetic acid (Ni-NTA) as a model ligand, we demonstrate that the attachment of lipid anchored Ni-NTA and PEG molecules to distinct lipid domains in nanoparticles can enhance liposome binding to cancer cells by increasing ligand clustering and reducing steric hindrance. We then use this technique to enhance the binding of RGD-modified liposomes, which can bind to integrins overexpressed on many cancer cells. These results demonstrate the potential of lipid phase separation to modulate the spatial presentation of targeting and shielding molecules on nanocarriers, offering a powerful tool to enhance the efficacy of NP drug delivery systems.

## 1. Introduction

The physicochemical properties of nanoparticles (NPs) play an important role in their therapeutic efficacy when used in drug delivery systems. A critical step in NP administration is interaction with a target cell surface. NPs are often engineered with two classes of surface modifications that affect this interaction: (1) targeting moieties to promote NP interactions with specific receptors and cell types,^1^ and (2) shielding polymers like polyethylene glycol (PEG) to increase *in vivo* circulation time and reduce the rate of in biological systems2. While many intrinsic properties of NPs affect ligand-receptor binding, such as stiffness,^3–5^ ligand mobility,^6–8^ and surface roughness,^9^ an important, often less explored feature is the spatial presentation of conjugated ligands on the NP surface. Ligand density is an important parameter that controls bond multivalency when targeting a NP to a cellular receptor.^10–13^ This density affects the overall binding strength of the NP to the target cell, or NP avidity, and ultimately contributes to the physiological fate of the NP and overall efficacy of drug delivery. As a result, optimizing the presentation of both targeting ligands and shielding polymers is important for modulating the NP avidity to the target cell receptor.^11^

Cells naturally modulate receptor avidity by dynamically rearranging membrane proteins and lipids in the cell membrane. Through the formation of lipid rafts, or highly ordered and cholesterol-rich regions of the cell membrane, cells can partition proteins into a lipid domain and exclude other proteins, which can control receptor density and function. This clustered arrangement of receptors facilitates not only receptor binding by significantly altering the binding kinetics, but also subsequent intracellular and intraorganelle signaling by lowering the activation threshold of receptors,^14, 15^ and it is a common feature of immune cell signaling.^16^ In addition, lipid rafts can exclude sterically hindering proteins and inhibitory signaling proteins that affect receptor binding and activation.^17^

Analogous to cellular membranes, the arrangement of ligands on a nanoparticle surface can also tune nanoparticle binding to target molecules. The clustered arrangement of ligands can enhance multivalent cooperativity^18–20^ by facilitating the rapid recapture of unbound or dissociating ligands by cell receptors. As a result, the enhanced binding of NPs to cells can lead to more efficient activation of cellular receptors through subsequent cluster-dependent signaling.^21–24^ One way to control ligand spatial presentation on nanoparticles is through lipid phase separation. By mixing saturated and unsaturated phospholipids with cholesterol, bilayer vesicles can undergo phase separation between the saturated liquid ordered (l_o_) and unsaturated liquid disordered (l_d_) phases, in a similar manner as cellular lipid rafts.^25^ Harnessing lipid phase separation to control ligand presentation on NPs has previously been used to enhance delivery of therapeutic cargo to cells ^26, 27^ and increase apoptotic signaling in target cells. Because many ligand-receptor binding systems are dependent on the surface concentration of the receptor, as well as the inter-ligand spacing on the targeting materials,^29, 30^ we wondered if lipid phase separation can be more broadly used to tune lipid nanoparticle avidity to cells for other types of binding interactions. Understanding the answer to this question will guide better design of NPs for targeted drug delivery.

In this study, we demonstrate an approach for assembling lipid bilayer-based NP vesicles that display both targeting ligands and shielding PEG polymers with varying surface densities. By varying composition and lipid chain saturation, we can assemble vesicles with varying sizes of membrane domains and spatially control localization of both ligands and PEG into distinct or colocalized domains. Using several phase-separated lipid compositions and ligands of varying binding strength, we identify key design principles for controlling and optimizing the avidity of vesicles to cells by controlling the location of targeting and sterically inhibiting molecules on the NP surface. We then apply these design principles to tune the performance of a therapeutically relevant ligand in cancer research: integrin-binding peptide sequence Arg-Gly-Asp (RGD), which is known to be dependent on inter-ligand spacing.^29, 31^ Our modular and readily accessible approach presents a powerful tool for using lipid phase separation to tune NP avidity without changing overall concentration of ligand-bound lipid or engineering new molecules with a different binding affinities. This technique can open the door to future technologies, including optimizing binding with lower concentration of ligand per particle, as well as stimuli responsive NPs that can harness phase separation to dynamically change their avidity to a target cell based on their environment.

## 2. Materials and Methods

### 2.1 Materials

Lipids including 1,2-dioleoyl-sn-glycero-3-phosphocholine (DOPC), 1,2-distearoyl-sn-glycero-3-phosphocholine (DSPC), cholesterol (Chol), 1,2-dioleoyl-sn-glycero-3-[(N-(5-amino-1-carboxypentyl)iminodiacetic acid)succinyl] (nickel salt) (DGS-NTA-Ni), 1,2-distearoyl-sn-glycero-3-phosphoethanolamine-N-[methoxy(polyethylene glycol)-2000] (18:0 PEG2000-PE), 1,2-dioleoyl-sn-glycero-3-phosphoethanolamine-N-[methoxy(polyethylene glycol)-2000] (ammonium salt) (18:1 PEG2000-PE), 1,2-dioleoyl-sn-glycero-3-phosphoethanolamine-N-(lissamine rhodamine B sulfonyl) (18:1 Rho), 1,2-dioleoyl-sn-glycero-3-phosphoethanolamine-N-(7-nitro-2-1,3-benzoxadiazol-4-yl) (ammonium salt) (18:1 NBD), 1,2-distearoyl-sn-glycero-3-phosphoethanolamine-N-[4-(p-(cysarginylglycylaspartate-maleimidomethyl)cyclohexane-carboxamide] (sodium salt) (DSPE-RGD), 1,2-dioleoyl-sn-glycero-3-phosphoethanolamine-N-[4-(p-(cysarginylglycylaspartate-maleimidomethyl)cyclohexane-carboxamide] (sodium salt) (DOPE-RGD) and 1,2-dioleoyl-sn-glycero-3-phosphoethanolamine-N-(cap biotinyl) (sodium salt) (18:1 Biotinyl Cap PE) were purchased from Avanti Polar Lipids. 1,2-dipalmitoyl-sn-glycero-3-phosphoethanolamine-N-(7-nitro-2-1,3-benzoxadiazol-4-yl) (16:0 NBD) were purchased from Invitrogen. Phosphate-buffered saline (PBS) tablets were obtained from Sigma-Aldrich. Cell media components, RPMI, fetal bovine serum (FBS) and penicillin/streptomycin were purchased from Thermo Fisher (Gibco). Streptavidin magnetic beads, calcein AM, and NucBlue Hoechst 33342 were purchased from Thermo Fisher. RGD peptide (GRGDNP) was purchased from MedChemExpress. Triton-X-100 (LabChem) was purchased from Fisher Scientific.

### 2.2 Large Unilamellar Vesicle Formation

Thin film hydration was used to prepare large unilamellar vesicles (LUVs). Lipid compositions for each vesicle formulation are provided in the Supporting Information Tables S1-S6. Lipid formulations in chloroform solution were dried down under a nitrogen stream to create a thin film on the bottom of a glass tube. Lipid films were placed in vacuum for 2 hours to remove excess chloroform. Films were rehydrated with 1x PBS (~290 mOsm) and incubated overnight at 60°C. Vesicles were vortexed and extruded using an Avanti mini extruder through a 100nm polycarbonate membrane for 7 passes. Vesicles containing saturated phospholipids were heated to 70°C on a hot plate during the extrusion process. Vesicles were characterized with regards to size and zeta potential by dynamic light scattering (DLS) using a Malvern Zetasizer Nano. Phase separation was quantified via a Förster resonance energy transfer (FRET) assay (see Supporting Experimental Section).

### 2.3 Cell Culture

Jurkat and U937 cells were obtained from the American Type Culture Collection (ATCC) without further authentication. K562 cells obtained from ATCC were gifted from the Leonard Lab (Northwestern University). Jurkat, U937, and K562 cells were cultured in RPMI 1640 supplemented with 10% FBS and 1% Penicillin-Streptomycin.

### 2.4 Vesicle-Cell Binding Studies

Vesicle binding to cells was characterized with flow cytometry. For dose response studies, assays were performed in a 96-well round bottom plate. For single concentration studies, assays were performed in Eppendorf tubes. Cells were suspended in flow buffer (PBS containing 1% FBS), and then seeded at a concentration of 100,000 cells per well/tube. Test vesicles were added to each well/tube at the specified concentrations and incubated with cells for 30 minutes at room temperature. Cells were washed 2 (tubes) to 3 (plates) times, where cells were spun down at room temperature in a centrifuge for 3 to 5 minutes until a pellet formed in each well/tube, and the supernatant was aspirated out and replaced with fresh flow buffer. Flow cytometry was performed on cells using a BD Fortessa LSRII using 550 nm excitation laser and 582/15 nm emission for rhodamine-labeled vesicles and 488 nm excitation laser 530/30 nm emission with 505 nm long pass filter for NBD-labeled vesicles. At least 5,000 events (for plate-based) to 10,000 (for tube-based) flow cytometry measurement events were selected by gating cell population based on forward vs. side scatter (FSC-A vs. SSC-A). Doublets were ruled out by gating based on FSC-A vs. FSC-H (Supporting Figure S1). Flow cytometry data including median fluorescence intensity (MFI) was analyzed using FlowJo software (Version 10.8.1). Fold change was calculated as the MFI of ligand-containing vesicle / MFI of unconjugated control vesicle within a replicate (paired) and the means of at least 3 fold changes was calculated.^32^

### 2.5 Vesicle-Bead Binding Assay

Vesicle binding to streptavidin beads was also characterized via flow cytometry. 2 μL of Pierce Streptavidin Magnetic Beads were added to PBS, and test vesicles were added to each tube at 500 uM lipid and incubated with beads in a 100 μL reaction for 30 minutes at room temperature. Beads were drawn down to the bottom of the solution with a magnet and the supernatant was replaced with fresh PBS, repeated three times. 10,000 flow cytometry measurement events were selected by gating bead population based on forward vs. side scatter (FSC-A vs. SSC-A), selecting for larger beads to rule out dust and debris. Doublets were eliminated by gating based on FSC-A vs. FSC-H (Supporting Figure S2). Flow cytometry data including median fluorescence intensity (MFI) was analyzed using FlowJo software.

### 2.6 Confocal Microscopy

Cells were stained with 500 ng/mL of calcein AM for 30 minutes, washed, and resuspended in full media. Vesicles were added to the cells at 500 μM final lipid concentration to 100,000 cells in 100 μL of 50% complete media, 50% PBS and incubated for 2 hours at 37°C for binding and internalization. After 2 hours, cells were washed 2x and resuspended in full media. NucBlue was added to the cells and imaged using confocal microscopy (Nikon Eclipse Ti).

### 2.7 Statistical Analysis

All significance tests, including 2-way ANOVA, multiple comparisons, and non-linear fits, were performed using Prism (GraphPad). For fold change data, all statistical analyses were conducted on the log-normal distribution [log(fold change)] in order to assume normal distribution.^32^

## 3. Results

### 3.1 Phase separation increases the avidity of a model weak-binding ligand

We prepared 100 nm vesicles with homogenously distributed and phase-separated lipids to probe how ligand density affects liposome-cell binding. Several different vesicle formulations were prepared with 0:1, 1:1, 2:1, 3:1 and 1:0 ratios of DSPC:DOPC, a saturated and unsaturated phospholipid, respectively, with a constant amount of cholesterol (30 mol%) and DSPE-PEG2000 (1 mol%) (Supporting Table S1). These formulations have been previously tested and characterized by our group using FRET analysis (See Supporting Experimental Section) to produce phase-separated vesicles. 0:1 and 1:0 vesicles containing only DOPC or DSPC and cholesterol have uniform distribution, and ternary vesicles containing DOPC, DSPC, and cholesterol show more phase separation as the ratio of DSPC:DOPC increases.^28^ To study whether phase separation can enhance vesicle avidity through ligand clustering, we chose to investigate Ni^2+^-nitrilotriacetic acid (Ni-NTA) as a model weak-binding ligand. Ni-NTA is a positively charged molecule commonly used in protein purification due to its interaction with polyhistidine tags, and has also been used to conjugate his-tag proteins to vesicles.^33–35^ Additionally, high Ni-NTA concentration on vesicles has been shown to increase nonspecific binding to cells,^36, 37^ which we suspect is due to charge-based interactions between Ni-NTA and histidine-rich proteins expressed on cell surfaces.^38, 39^ When added to the vesicle formulation, DGS-NTA(Ni), an unsaturated phospholipid covalently bound to Ni-NTA, should localize to the smaller, unsaturated DOPC domain resulting in a higher local density of DGS-NTA(Ni) on the vesicle surface, which we hypothesized would increase vesicle avidity (Figure 1a). Because 1 mol% Ni-NTA did not increase nonspecific binding to cells when uniformly present on DOPC vesicles, we chose this condition to study the effect of increasing the local density of DGS-NTA(Ni) through phase separation on cell binding (Supporting Figure S3).

**Figure 1.**
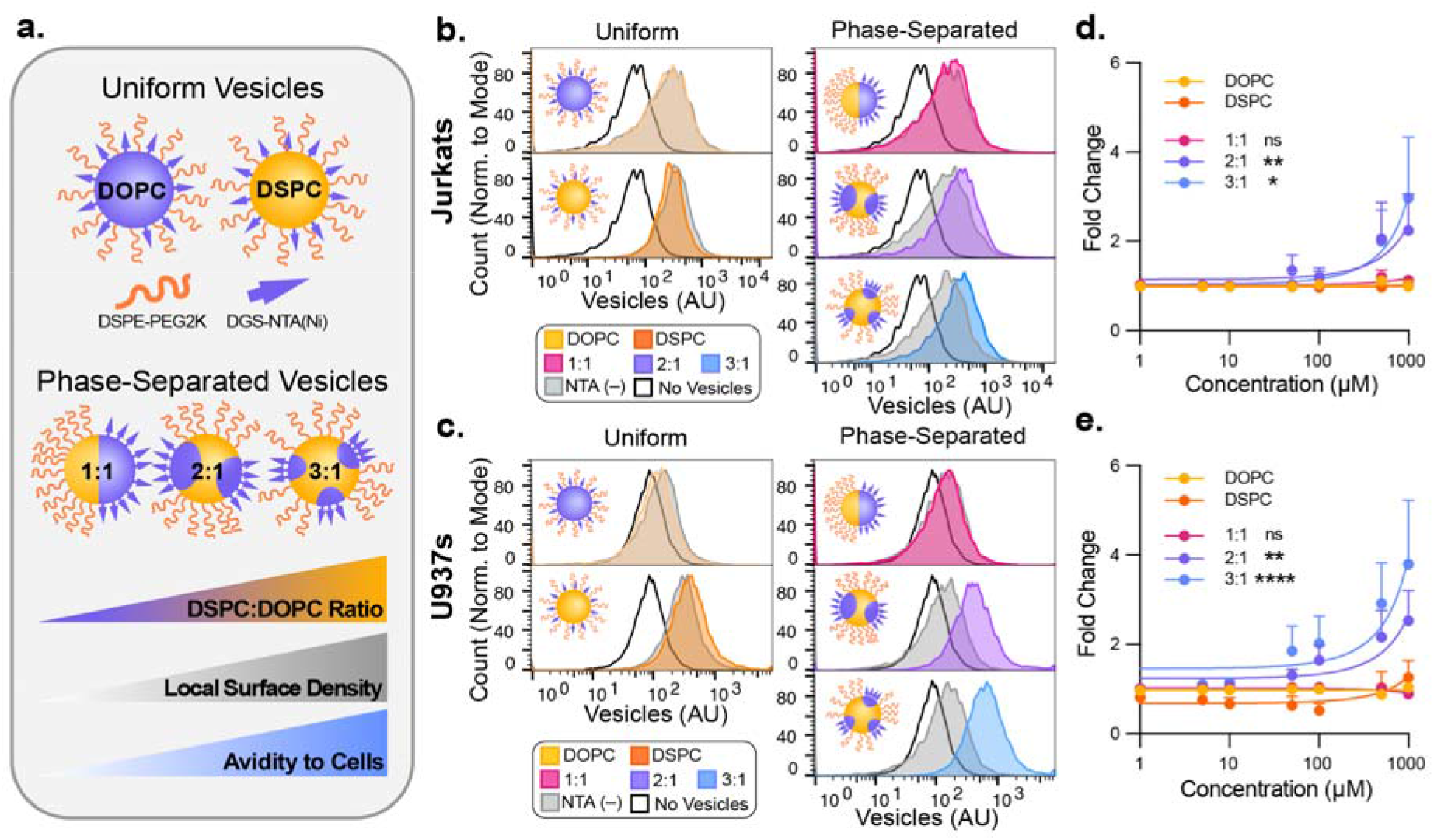
Phase separation enhances Ni-NTA vesicle avidity to cells. (a) Schematic of the vesicle compositions used in these experiments. By varying the ratio of DSPC:DOPC, local surface density of DGS-NTA(Ni) can be increased, which increases avidity to cells. (b,c) Flow cytometry histograms of vesicle avidity to (b) Jurkat and (c) U937 cells at 500 μM. Black line plot represents unstained cells, grey plot represents unconjugated vesicles, and colored plots represent NTA-conjugated vesicles. (d,e) Fold change of median fluorescence intensity of (d) Jurkat and (e) U937 cells incubated with Ni-NTA vesicles over no-NTA control vesicles. Error bars represent one-sided SEM for three biological replicates conducted on different days (n=3). Significance testing represents results of two-way ANOVA over concentration and compositions with Dunnett’s multiple comparison tests with respect to DOPC. * p<0.05, ** p<0.01, **** p<0.0001.

To study vesicle avidity to cells due to phase separation-mediated Ni-NTA clustering, we incubated fluorescent phase-separated vesicles with and without 1 mol% DGS-NTA(Ni) with Jurkat and U937 cells, which are human T-cell and monocyte lines, respectively. Vesicle binding was analyzed via flow cytometry, and median fluorescence intensity (MFI) values for cells incubated with DGS-NTA(Ni) vesicles were normalized to background MFI values from corresponding cells incubated with non-functionalized vesicles to study the effects of DGS-NTA(Ni) clustering while ruling out any differences in avidity due to lipid composition.^40^ We observed that 2:1 and 3:1 DSPC:DOPC phase-separated vesicles, the compositions we expect to have the highest surface density of Ni-NTA ligands, enhanced DGS-NTA(Ni)-mediated cell binding. However, DGS-NTA(Ni) did not increase binding when added to 1:1 DSPC:DOPC and uniform DOPC vesicles (Figure 1b and 1c). We also tested vesicle binding to U937 cells and observed similar trends, indicating that enhancing DGS-NTA(Ni)-mediated cell binding via phase separation is not a phenomenon limited to one cell type (Figure 1d and 1e). Avidity effects were prominent at concentrations of 500 μM lipid vesicles and higher, and therefore 500 μM was chosen for future studies. We hypothesize that by increasing the ratio of DSPC:DOPC, we increased the local concentration of Ni-NTA on the vesicle surface in a similar manner to when we increased the global concentration of Ni-NTA in vesicles (Supporting Figure S3). Interestingly, 1:1 DSPC:DOPC did not increase Ni-NTA binding relative to DOPC vesicles, indicating a certain threshold of ligand density is required to observe enhanced avidity. Together, this data demonstrates that lipid phase separation provides a route to enhance vesicle-to-cell avidity for a low affinity, charge based interaction by increasing the clustering of DGS-NTA(Ni) lipids, and a critical threshold of phase separation is required to observe enhanced binding.

### 3.2 Phase separation can be used to modulate the steric effects of PEG on a model weak-binding ligand

Next, we investigated how modulating the location of PEG on NTA(Ni)-containing vesicles affects binding to cells. Besides improving nanoparticle stability and reducing biofouling, PEG molecules can undesirably shield ligands through steric effects.^41^ Cleavable PEG systems have been developed to take advantage of this effect by better exposing ligands after encountering a disease-specific stimuli.^42, 43^ Motivated by this shielding capacity, we hypothesized that segregation of PEG away from the ligand on the vesicle surface could improve ligand accessibility. To study the effect of PEG segregation on binding, we created Ni-NTA vesicles and incorporated varying ratios of unsaturated and saturated PEG lipids (DOPE-PEG and DSPE-PEG, respectively) at an overall constant PEG concentration of 1 mol% (Figure 2a). We added varying ratios of PEG lipids into both phase-separated Ni-NTA vesicles (3:1 DSPC:DOPC) and uniform vesicles (DOPC) and conducted binding studies (see Supporting Table S2 for lipid composition). In uniform vesicles, vesicle binding to cells was similar for all DOPE-PEG:DSPE-PEG ratios, consistent with our hypothesis that PEG lipids are evenly distributed across the vesicle surface (Figure 2b and Figure 2c). As anticipated, in phase-separated vesicles, the presence of DOPE-PEG in the same phase as the Ni-NTA groups decreased vesicle avidity and replacing DOPE-PEG with DSPE-PEG to sequester the PEG molecules in a separate domain from the Ni-NTA groups enhanced vesicle binding (Figure 2b and Figure 2c). This result suggests the behavior we observed with phase segregated vesicles is not due to chemical differences between the two types of lipid-modified PEG that were used. Because the addition of PEG-modified lipid can dissociate lipid domains,^44^ we conducted FRET studies to confirm that lipid domains existed at all PEG ratios (Supporting Figure S4, Supporting Table S5). Taken together, our results demonstrate that by simply tuning the location of PEG, we can tune phase-separated vesicle avidity to target cells, independent of the ligand’s binding affinity. Modifying the location of PEG or other shielding polymers away from targeting ligands could be used in lieu of ligand engineering techniques to tune overall nanoparticle avidity.

**Figure 2.**
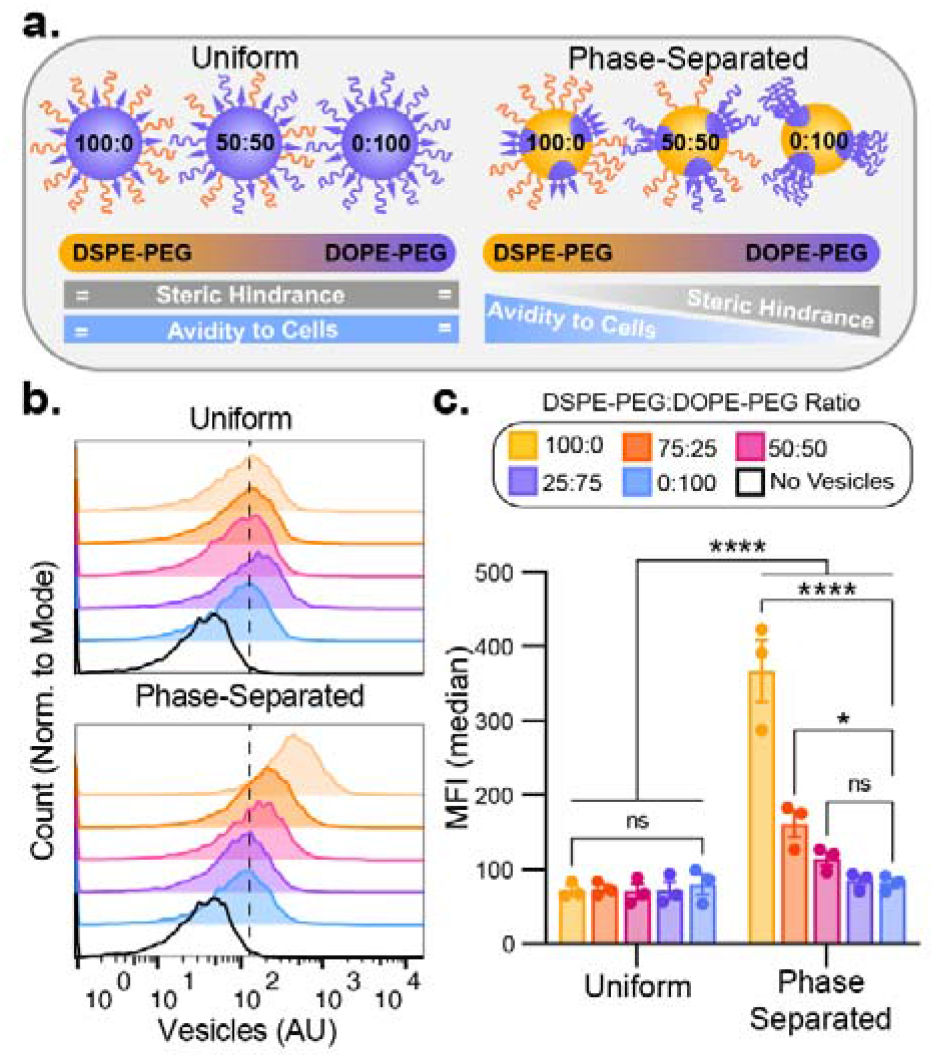
Vesicle avidity can be tuned by controlling the colocalization of PEG and Ni-NTA lipid. (a) Schematic depicting some of the vesicle compositions examined in this experiment. In uniform vesicles, changing the ratio of DSPE-PEG to DOPE-PEG does not affect the spatial distribution of PEG and does not affect vesicle avidity. In phase-separated vesicles, controlling this ratio modulates the degree of colocalization which can increase or decrease binding. (b) Flow cytometry histograms of uniform and phase-separated vesicle binding to Jurkat cells with varying ratios of DSPE-PEG:DOPE-PEG. Dotted line represents approximated median of uniform vesicle binding to cells. (c) Quantitative and statistical analysis of the flow cytometry data between phase-separated vs. uniform vesicles and within each type of vesicle with varying ratios of PEG lipids. Errors bars represent SEM for n=3. Significance testing represents the results of two-way ANOVA using Tukey’s multiple comparison tests. **** p < 0.0001, * p < 0.05.

### 3.3 Phase separation has a weaker effect on avidity in a model strong-binding ligand

After studying the effect of lipid phase separation and PEG location on the avidity of a weak-binding, vesicle-bound ligand, we set out to investigate how these phenomena compared with a strong-binding ligand. We used biotin-avidin interactions for these studies by measuring binding of biotinylated vesicles to streptavidin-coated magnetic beads. Uniform and phase-separated vesicles with 0.1 mol% unsaturated, biotinylated lipid (18:1 Biotinyl Cap PE) with varying ratios of DSPE-to DOPE-bound PEG were incubated with avidin coated magnetic beads, and binding was measured via flow cytometry (see Supporting Table S3 for lipid compositions). Phase-separated biotinylated vesicle binding to avidin beads was enhanced when compared to uniform vesicles, similar to the effect observed in most Ni-NTA vesicles (Figure 3). These results indicate that clustering through phase separation may enhance many different ligand-receptor systems across a wide range of binding affinities. Contrary to what we found with Ni-NTA, however, the ratio of DOPE-PEG to DSPE-PEG did not have a significant effect on the binding of biotin vesicles to avidin beads (Figure 3). One explanation for this is that the strength of the biotin-avidin interaction may be strong enough to overpower the steric effects of PEG. However, high molecular weight PEG has also been shown to have steric hindrance effects on biotin-avidin systems in some cases, including on biotinylated vesicles,^45, 46^ suggesting PEG location relative to targeting ligands may still modulate stronger binding protein interactions. Another possibility is that beads present targets statically, while cell membranes are fluid, which could explain the differences we see on binding.

**Figure 3.**
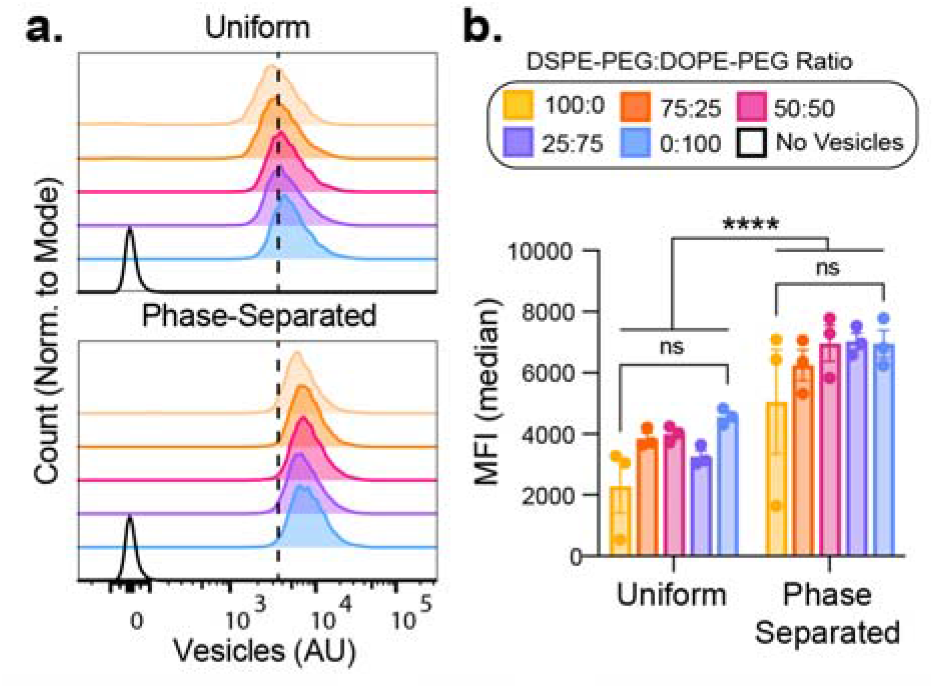
Steric hindrance-modulated vesicle avidity is less effective with strong binders. (a) Flow cytometry histograms of uniform and phase-separated vesicle binding to streptavidin beads with vesicles containing varying ratios of DSPE-PEG:DOPE-PEG. Dotted line represents approximate median binding of uniform vesicles to beads. (b) Quantitative and statistical analysis of the flow cytometry data between phase-separated vs. uniform vesicles and within each type of vesicle with varying ratios of PEG lipids. Error bars represent SEM for n=3. Significance testing represents the results of two-way ANOVA using Tukey’s multiple comparison tests.

### 3.4 Phase separation can enhance binding of vesicles with a therapeutically relevant ligand, RGD

To demonstrate translational utility of this technology, we sought to confirm that phase separation can enhance cell-binding of vesicles functionalized with a therapeutically relevant ligand. The Arg-Gly-Asp (RGD) peptide motif interacts with α_5_β_1_ and α_v_β_3_ integrins, which are overexpressed on many cancers, and therefore RGD-functionalized liposomes have been studied extensively for drug delivery.^47^ While the cyclic form of RGD is typically preferred due to its higher affinity to integrin and stability, the linear form is more readily available conjugated to lipid. Furthermore, a recent study demonstrated giant unilamellar vesicles functionalized with linear RGD were able to bind Jurkat cells.^48^ We therefore wanted to test whether phase-separated vesicles functionalized with RGD would enhance RGD-mediated binding (Figure 4a).

**Figure 4.**
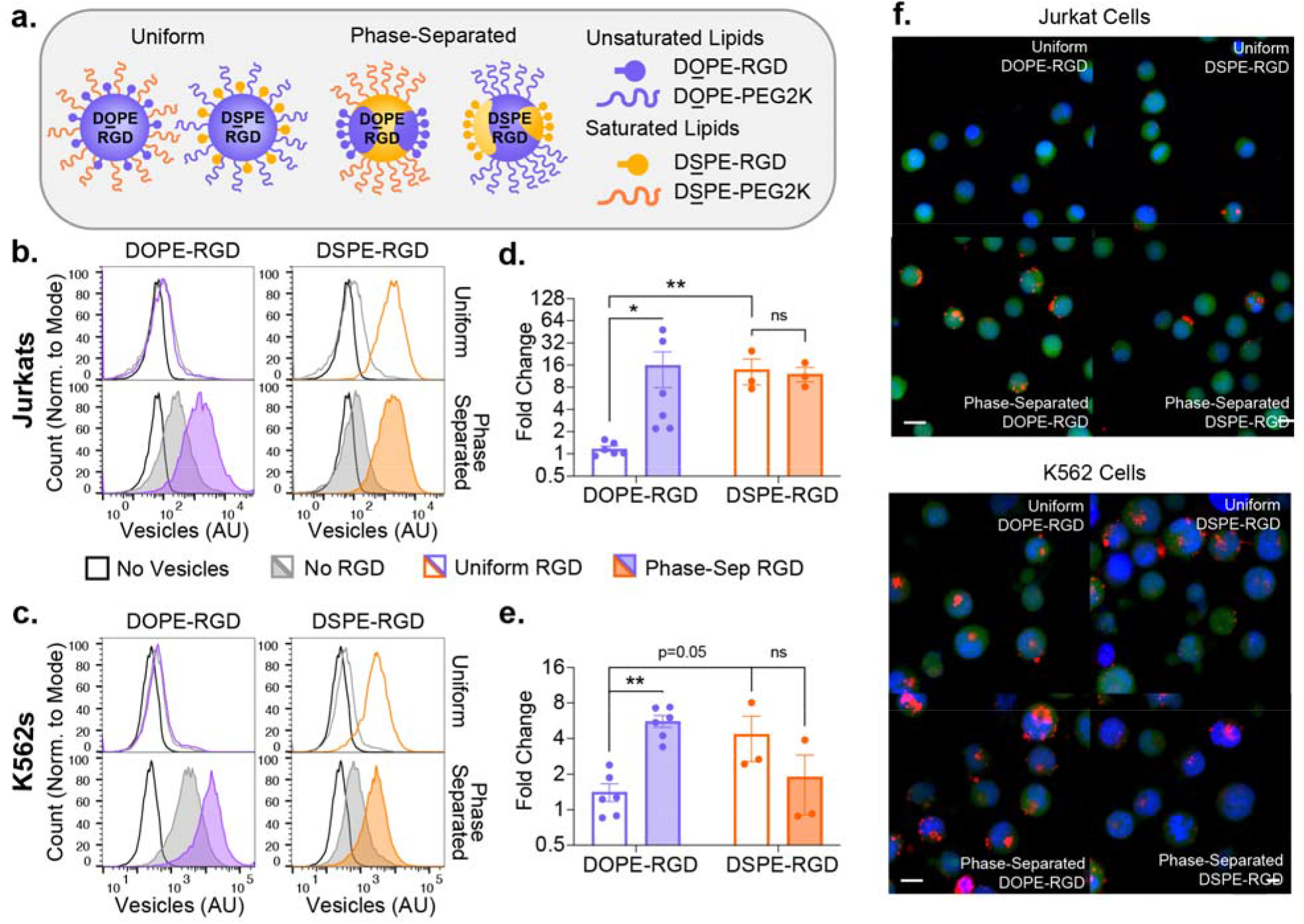
Phase separation can enhance RGD-mediated cell binding to Jurkat and K562 cells, but only when RGD is in the l_d_ phase and not the l_o_ phase. (a) Schematic depicting the vesicle compositions explored in this experiment. In uniform DOPC vesicles, the lipid anchor on the PEG and the RGD does not affect their local surface density. In phase-separated vesicles, RGD can be localized to either the l_o_ phase (yellow) or the l_d_ phase (purple), depending on the lipid anchor. (b,c) Flow cytometry histograms of uniform and phase-separated DOPE- and DSPE-RGD vesicle binding to (b) Jurkat and (c) K562 cells, as well as the no RGD controls. (d,e) Quantitative analysis of fold increase in MFI of (d) Jurkat and (e) K562 cells incubated with RGD vesicles over no RGD control vesicles. Error bars represent SEM for three (DSPE-RGD vesicles) to six (DOPE-RGD vesicles) measurements. Significance testing represents the results of two-way ANOVA with Tukey’s multiple comparison tests. * p<0.05, ** p<0.01. (f) Confocal microscopy images of Jurkat and K562 cells treated with uniform and phase-separated DOPE- and DSPE-RGD vesicles. Cells are stained with the nuclear dye NucBlue (blue), the viability dye calcein AM (green). Vesicles are doped with 0.1 mol% 18:1 Liss Rhod PE (red). Scale bar = 10 μm.

We created uniform and phase-separated vesicles with 10 mol% DOPE-RGD, confirmed phase separation using FRET (Supporting Figure S5), and tested binding to Jurkat and K562 cells, cell lines that both highly express the RGD-binding integrin α_5_β_1_.^49^ In uniform vesicles, 10 mol% RGD vesicles exhibited minimal binding on both Jurkat and K562 cells compared to unfunctionalized control vesicles (Figure 4b–e). When RGD was concentrated into lipid domains, binding was greatly enhanced to Jurkats (~15 fold) and K562s (~5 fold) (Figure 4b–e). To verify vesicle binding was due to specific interaction between vesicle-bound RGD and cellular receptors, we added a soluble RGD peptide that competitively binds to integrins. The addition of the soluble RGD peptides significantly reduced phase-separated vesicle binding, supporting the specificity of the vesicle-to-cell binding interaction (Supporting Figure S6). Interestingly, we did not see vesicle binding to Jurkat cells when RGD was uniformly distributed on the vesicle surface as previously reported, even at 10 mol% RGD.^48^ The discrepancy with previous data could be due to differences in vesicle size as the larger vesicles (2 μm diameter) used in the previous study have more RGD ligands and a larger surface for multivalent binding. In summary, phase separation enhances RGD-mediated vesicle binding, and supports the applicability of phase separation as a tool to enhance interactions of nanoparticle-bound, therapeutically relevant ligands with target cells.

### 3.5 The lipid saturation of RGD conjugation affects cell binding

Finally, we investigated whether clustering RGD lipids in the l_o_ phase could also enhance RGD-mediated cell binding. In our previous experiments, the ligand was conjugated to an unsaturated lipid, and therefore the ligand should localized into the ld phase during phase separation. Therefore, we wanted to compare how concentrating the ligand into the l_o_ phase affected vesicle avidity compared to the l_d_ phase, since the fluidity of the local membrane affects the mobility of the ligand and therefore has the potential to affect vesicle binding (Figure 4a).^6–9^ To investigate this, we created uniform and phase-separated vesicles with DSPE-RGD, which should localize into the l_o_ phases of vesicles (Figure 4a and Supporting Table S4 for full composition). We observed that uniform vesicles containing DSPE-RGD and DOPE-RGD differentially bound to cells. In particular, uniform vesicles containing DSPE-RGD bound significantly stronger to Jurkat cells compared to uniform DOPE-RGD vesicles (Figure 4b, c), while in K562 cells this effect was less pronounced (p=0.0503, Figure 4d, e). The differences in integrin expression between these cells could play a role in the differences in binding, as integrins might have their unique optimal special presentation.^49^ Additionally, phase-separated vesicles containing DSPE-RGD did not bind stronger than uniform DSPE-RGD vesicles in either cell type. FRET analysis to confirm lipid domains demonstrated that DSPE-RGD phase-separated vesicles did exhibit domain formation (Figure S5). However, this composition also exhibited a lower melting temperature compared to phase-separated DOPE-RGD, which could indicate a morphological or chemical stability difference between domains that might affect binding. Furthermore, it is possible that DSPE-RGD segregates into small clusters that are undetectable by FRET even in compositions that should lead to uniform DSPE-RGD distribution, which is supported by our previous studies.^28^ Headgroups also affect domain localization and could also affect how well RGD lipids phase separate, as seen similarly with fluorophores.^50^ Regardless, our results demonstrate that using phase-separation to increase avidity of DSPE-RGD is not effective in contrast to DOPE-RGD vesicles, suggesting that the differences between saturation of ligand-bound lipid and local lipid environment may have a profound effect on binding.

We also studied RGD vesicle interactions with cells via confocal microscopy. Jurkat and K562 cells were incubated with the same RGD vesicle compositions mentioned above for 2 hours (red), and stained with a nuclear dye (NucBlue, blue). Images on the confocal microscope confirmed similar trends to those observed with flow cytometry, where phase separation increased binding of DOPE-RGD vesicles but not DSPE-RGD vesicles in both cell types (Figure 4f and Supporting Figure S7). Moreover, our images demonstrate improved interactions of DOPE-RGD phase-separated vesicles compared to their uniform controls, suggesting that phase separation of ligands on nanoparticles to improve vesicle avidity to a target cell may also have the potential to enhance delivery of therapeutic cargo. Further studies will be required to investigate how phase separation affects vesicle internalization.

## 4. Discussion

The observed importance of membrane organization motivates exploration of how similar phenomena can be mimicked in synthetic systems, which can not only improve our understanding of these processes but also be applied to new technologies. Cells spatially control their membrane because some receptors have optimal distances to induce signaling,^23^ but overexpression of receptors which would increase receptor proximity can lead to disease through constitutive activation.^51^ To overcome the need to use gene expression changes—a slow process—to induce changes in signaling kinetics, cells utilize their membranes to rapidly reorganize and cluster receptors together in response to certain cell states or environmental conditions so that they are better poised to initiate intracellular and intraorganelle signaling and reduce the effects of noise.^15, 52^ Furthermore, clustering can also affect avidity by increasing ligand rebinding, and ultimately the probability of a binding event initiating signal transduction.^53, 54^ This is observed in T cells, which need to balance receptor affinity and expression in order to ultimately optimize their overall functional avidity.^55, 56^ To do that, T cells utilize receptor clustering to dynamically change their avidity,^54^ which has also been harnessed in CAR-T cell engineering as well.^57, 58^ Clustering of low affinity ligands or receptors to change the overall avidity of ligand-receptor interactions is therefore a potent, biologically relevant strategy to not only prevent off-target signaling but also dynamically enhance signaling in response to environmental changes.

We believe these same cellular principles of modulating cellular avidity through clustering of lower affinity receptors can also be useful when designing therapeutic nanoparticles. Our group has recently demonstrated that clustering of receptors on vesicles using phase separation can be significant in inducing intracellular signaling.^28^ The current work demonstrates that phase separation can enhance interactions between a wider range of biologically relevant, low affinity interactions, and furthermore be harnessed to enhance binding avidity of vesicles by changing the location of sterically shielding molecules. Together, our results present a route to use smaller amounts of targeting or bioactive ligands while having a similar effect on target cell binding. Because many therapeutic ligands are expensive to produce, we believe phase separation in nanoparticles is a readily achievable route to enhance local ligand concentration and require overall less ligand per particle. Using both phase separation and modulating ligand expression, we now have two tools that can be used to fine-tune nanoparticle avidity. Furthermore, phase separation uses less surface area on a nanoparticle, and therefore allows for building unique nanoparticles with multiple ligands with controlled spatial orientation. We envision that this work is another tool that can be used to expand on vesicle technologies with multimodal functionalities.^59^

Beyond mimicking the cell membrane, phase separation could also be used to engineer stimuli-responsive vesicles to enhance drug delivery. Phase separation can be induced through multiple stimuli including tension,^60^ pH,^61^ temperature,^62^ small molecules,^63^ peptides,^64^ proteins,^65,66^ and association with soluble oligomerizing molecules.^67, 68^ Phase-separated vesicles could be used in conjunction with these stimuli to dynamically change the presentation of nanoparticle-bound ligands. For example, with a temperature responsive vesicle,^62^ temperature changes could be used to change ligand presentation, cell binding and release of encapsulated cargo. Furthermore, PEG phase separation could be used to dynamically modulate steric shielding events in response to environmental conditions of the nanoparticle. By using lipid phase separation, new classes of stimuli-responsive nanoparticles can be developed for enhancing drug delivery by dynamically controlling the presentation of both targeting and shielding molecules.

## 5. Conclusion

In this work, we demonstrated that lipid phase separation can be used to spatially control ligands on lipid vesicles, which can be used to modulate the overall avidity of a NP to a target cell. Phase separation can enhance NP binding to cells by both enhancing ligand binding through clustering and adjusting the steric hindrance of PEG molecules. This method is readily adoptable and accessible to anyone working with vesicles. We also demonstrate that the saturation of the lipid anchor can ultimately affect ligand-cell binding and should also be considered when creating lipid-based nanoparticles. Future work will look at using phase separation to spatially pattern multiple ligands, and the extent to which phase separation can enhance multifunctional lipid-based NPs. Similar to how cells rearrange their membranes and ligands to activate receptor signaling, we expect nanoparticles that mimic this feature should better bind and activate target cells, which should be useful for applications like drug delivery.

## Supporting Information

Supporting experimental section, binding of vesicles as a function of NTA concentration, FRET analysis of vesicle phase separation, RGD blocking, enlarged microscopy, and vesicle compositions and characterization are included in the Supporting Information.

## Corresponding Author

**Neha P. Kamat -** Department of Biomedical Engineering, McCormick School of Engineering and Center for Synthetic Biology, Northwestern University, Evanston, IL, 60208, United States;

## Authors

**Timothy Q. Vu -** Department of Biomedical Engineering, McCormick School of Engineering, Northwestern University, Evanston, Illinois 60208, United States

**Lucas E. Sant’Anna -** Department of Biomedical Engineering, McCormick School of Engineering, Northwestern University, Evanston, Illinois 60208, United States

## Author Contributions

L.E.S. and T.Q.V. contributed equally to this work. L.E.S., T.Q.V., and N.P.K. conceived the study and designed experiments. L.E.S. and T.Q.V. performed the experiments. L.E.S., T.Q.V. and N.P.K. analyzed the data. All the authors contributed to writing the manuscript.

## Acknowledgements

L.E.S. was supported by Northwestern’s Academic-Year Undergraduate Research Grant and the Jaharis Family Foundation through the Michael Jaharis Undergraduate Research Fellowship.

T.Q.V. was supported by the National Institutes of Health Training Grant (T32GM008449) through Northwestern University’s Biotechnology Training Program. This work made use of the Northwestern RHLCCC Flow Cytometry Facility (NCI CA060553) and Northwestern NUANCE center (NSF ECCS-2025633 & NSF DMR-1720139). We thank Kamat lab members for helpful discussion.

## Notes

N.P.K and T.V. are inventors on a U.S. provisional patent submitted by Northwestern University that covers material encompassing this study on organizing proteins in lipid membranes via phase segregation.

