## Supporting Information for "Enhancing functionalized liposome avidity to cells via lipid phase separation"

**TABLE OF CONTENTS**

**Supporting Experimental Section** ………………………………………………………… 2

**Supporting Figures** ………………………………………………………………………… 3

**Figure S1:** Gating strategy for cell-based flow cytometry assays.

**Figure S2:** Gating strategy for bead-based flow cytometry assays.

**Figure S3:** Vesicle-Jurkat binding at varying concentrations of DGS-NTA(Ni).

**Figure S4:** FRET study of DGS-NTA(Ni) vesicles.

**Figure S5:** FRET study of RGD vesicles.

**Figure S6:** RGD block assay.

**Figure S7:** Enlarged microscopy images of RGD vesicle uptake

**Supporting Tables** …………………………………………………….…………………… 10

**Table S1:** Vesicle compositions and DLS data for Figure 1.

**Table S2:** Vesicle compositions and DLS data for Figure 2.

**Table S3:** Vesicle compositions for Figure 3.

**Table S4:** Vesicle compositions and DLS data for Figure 4.

**Table S5:** Vesicle compositions for Figure S2.

**Table S6:** Vesicle compositions for Figure S3.

**SUPPORTING EXPERIMENTAL SECTION**

*Fluorescence Resonance Energy Transfer (FRET) Assays*

Extent of lipid mixing was measured by FRET assay. Vesicles containing 0.5 mol% of 18:1 Rhodamine (acceptor) and 0.5 mol% 16:0 NBD (donor) lipid dye were added to a cuvette and their fluorescence was measured at 460 nm excitation and 535 nm and 583 nm emission using an Agilent Cary Eclipse Fluorimeter. After measurements were taken, vesicles were lysed with 0.1% Triton-X to read the unquenched fluorescence for normalization. For temperature ramp experiments, fluorescence was measured from 25°C to 58°C at 1°C steps. Normalized FRET ratio corresponds to extent of phase-separation, and is defined as:

N𝑜𝑟𝑚𝑎𝑙𝑖𝑧𝑒𝑑 𝐹𝑅𝐸𝑇 𝑟𝑎𝑡𝑖𝑜 = $\frac{F_{donor}/F_{acceptor}}{F_{donor,triton}/F_{acceptor,triton}}$

*RGD Peptide Block Assay*

RGD peptide block assays were performed similar to vesicle-cell binding assays, except before the addition of vesicles to cell culture, soluble RGD peptide was combined with cells suspended in PBS at 500 μM (10x RGD mol% of RGD vesicles) and allowed to bind for 30 minutes. Cells were then spun down and supernatant was aspirated to remove unbound RGD, and cells were resuspended in fresh flow buffer. Binding assay from the experimental section was subsequently performed.

**SUPPORTING FIGURES**

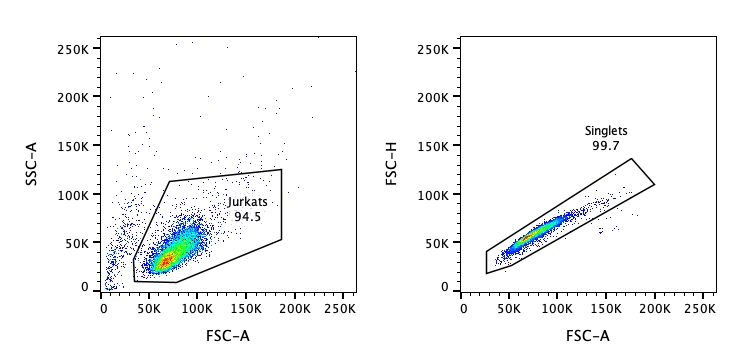

**Figure S1.** Gating strategy for cell-based flow cytometry assays. Cells were first gated on SSC-A vs. FSC-A. Then, doublets were ruled out by gating FSC-H vs FSC-A.

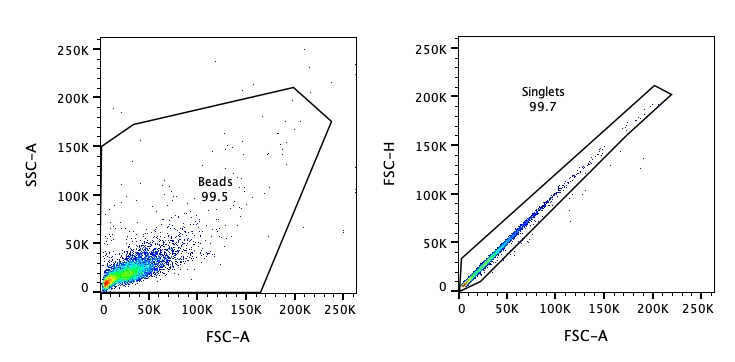

**Figure S2.** Gating strategy for bead-based flow cytometry assays. Beads were first gated on SSC-A vs. FSC-A. Then, doublets were ruled out by gating FSC-H vs FSC-A.

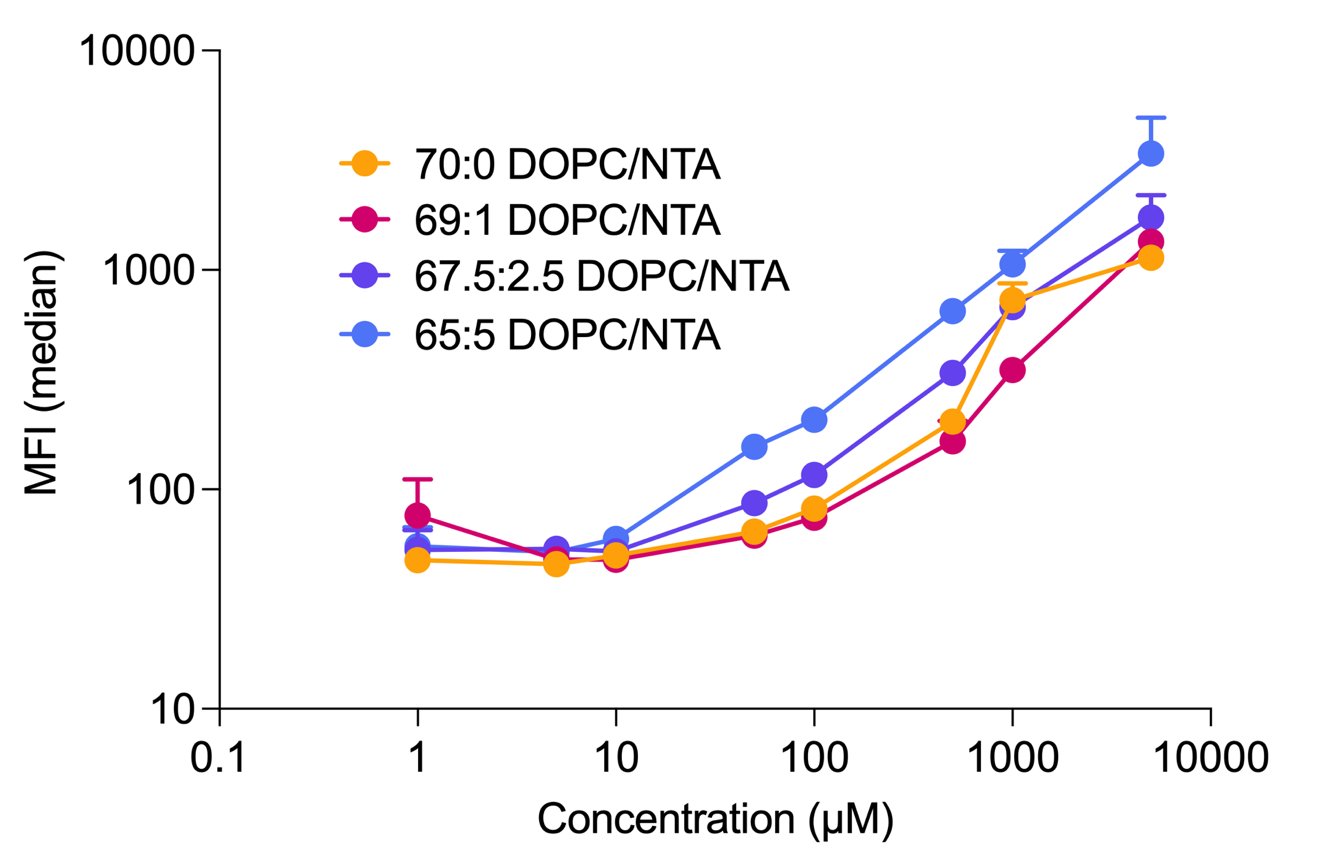

**Figure S3.** Binding curves for vesicles containing varying concentrations of DGS-NTA(Ni) lipid. 1 mol% DGS-NTA(Ni) was chosen for future experiments due to minimal non-specific binding at 500 µM. Error bars represent SEM from n=2.

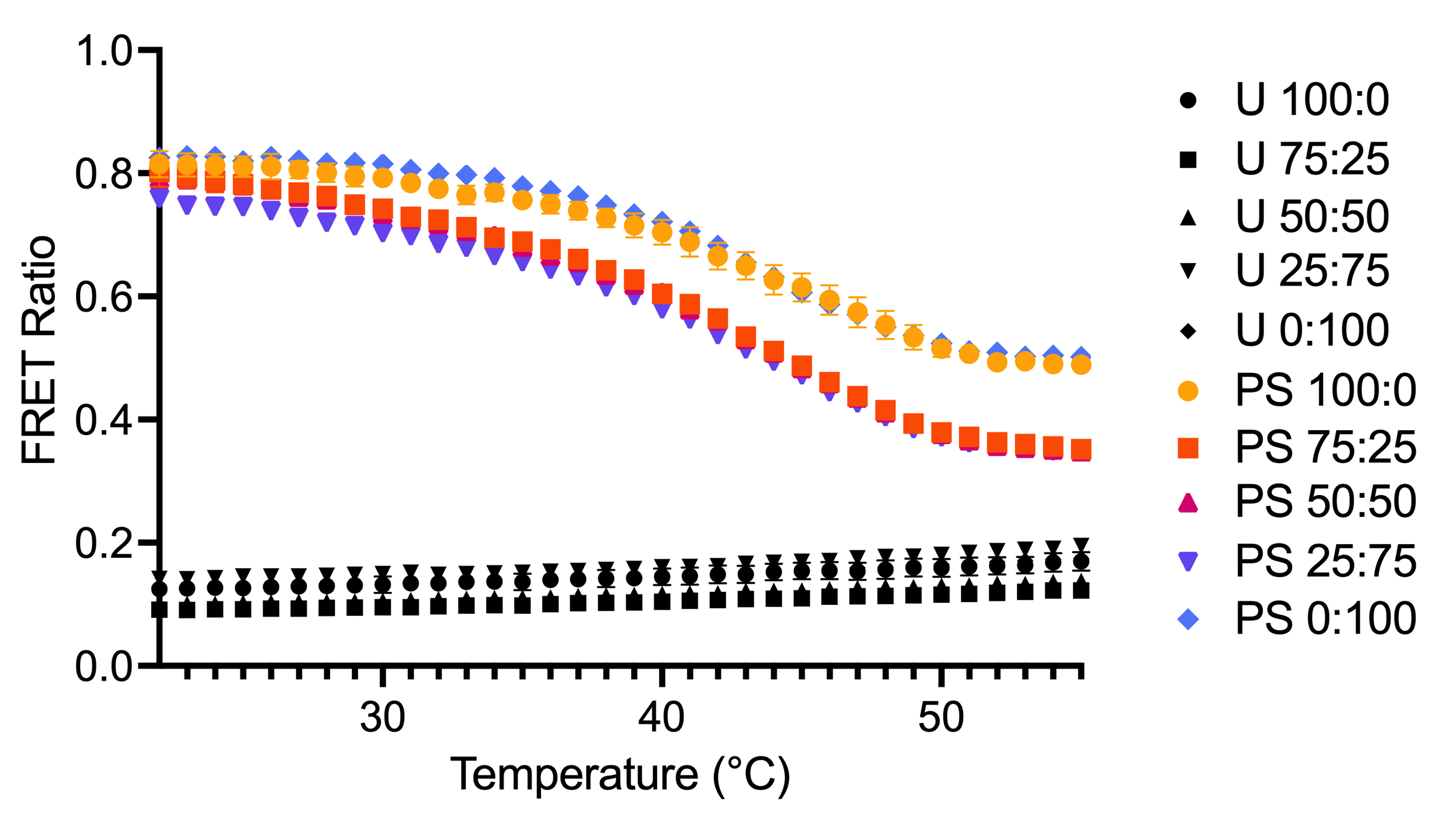
 **Figure S4.** FRET temperature ramps for DGS-NTA(Ni) containing phase-separated (PS) and uniform (U) vesicles. Error bars represent SEM from three measurements across two different vesicle preparations. Vesicles exhibiting a higher FRET ratio are more phase-separated. PS vesicles indeed exhibit more phase-separation at room temperature, and FRET ratios approach that of uniform vesicles upon heating due to temperature induced dissolution of domains. PEG content does not disrupt domain formation in PS vesicles, and trends in minor differences are not reflected in binding avidity assays.

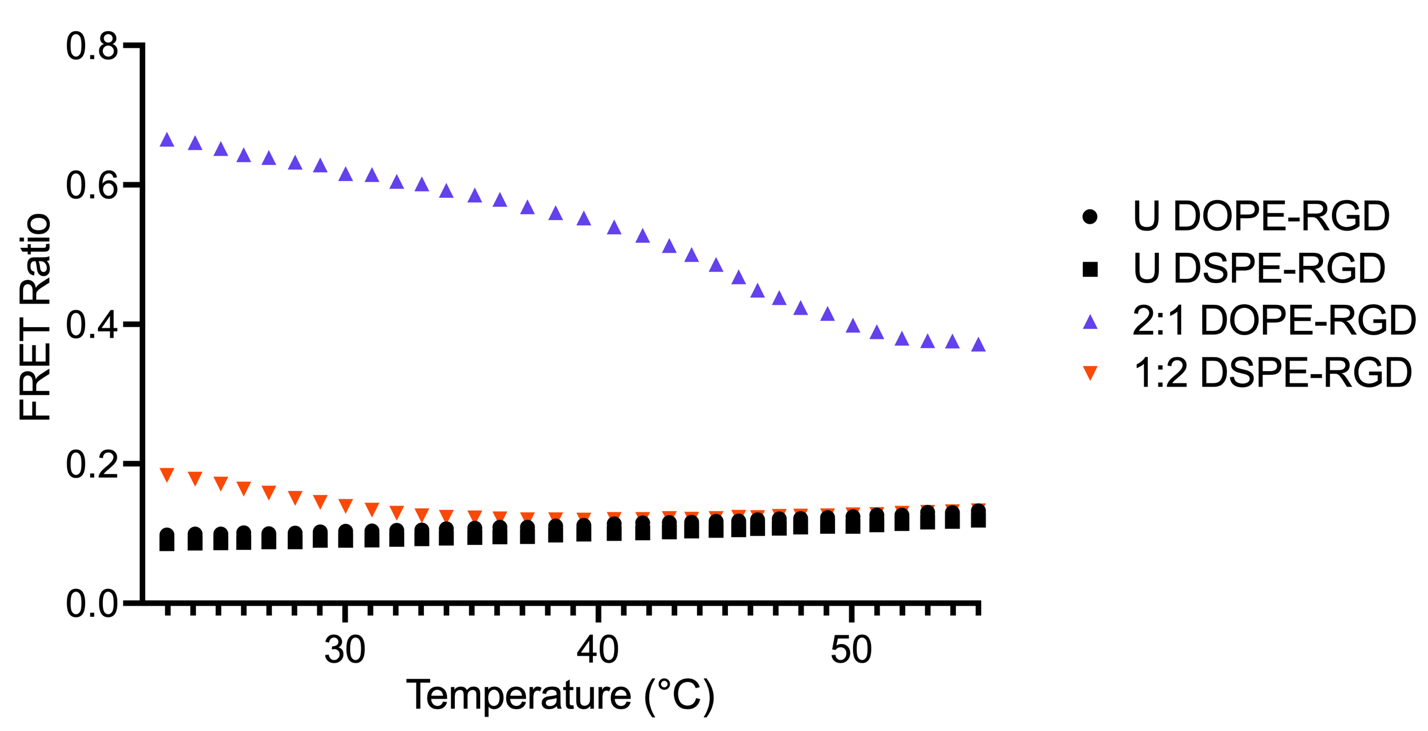

**Figure S5.** FRET temperature ramps for DOPE-RGD and DSPE-RGD containing phase-separated and uniform vesicles. Error bars represent SEM from three measurements across two different vesicle preparations (not visible behind symbols). Both 2:1 DOPE-RGD and 1:2 DSPE-RGD vesicle exhibit phase-separation at room temperature, although 1:2 DSPE-RGD vesicles to a lesser extent. Additionally, 1:2 vesicle domains dissolve at a lower temperature than 2:1 vesicles.

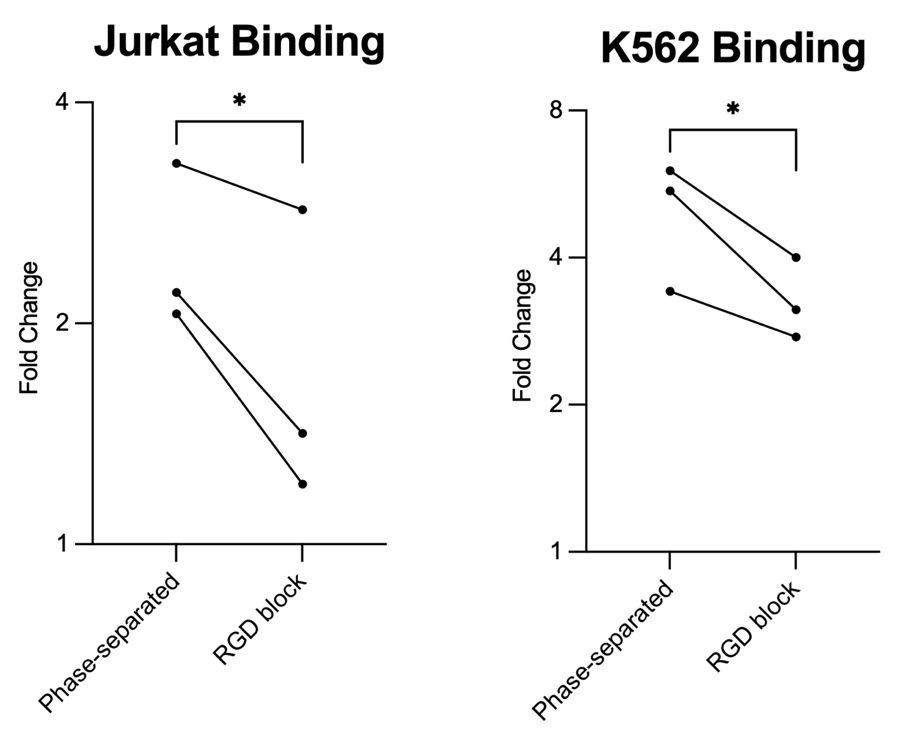

**Figure S6.** RGD peptide block assay. Addition of RGD peptide to vesicle-cell binding assay caused a significant decrease in the vesicle binding to both Jurkat and K562 cells. However, blinding was not completed blocked by RGD peptide. Multivalency of RGD-functionalized vesicles may account for this effect by displacing bound soluble RGD peptides, but nevertheless a significant reduction in binding verifies the specificity of RGD vesicles. * = p<0.05. One way paired t-test was performed on log(fold change).

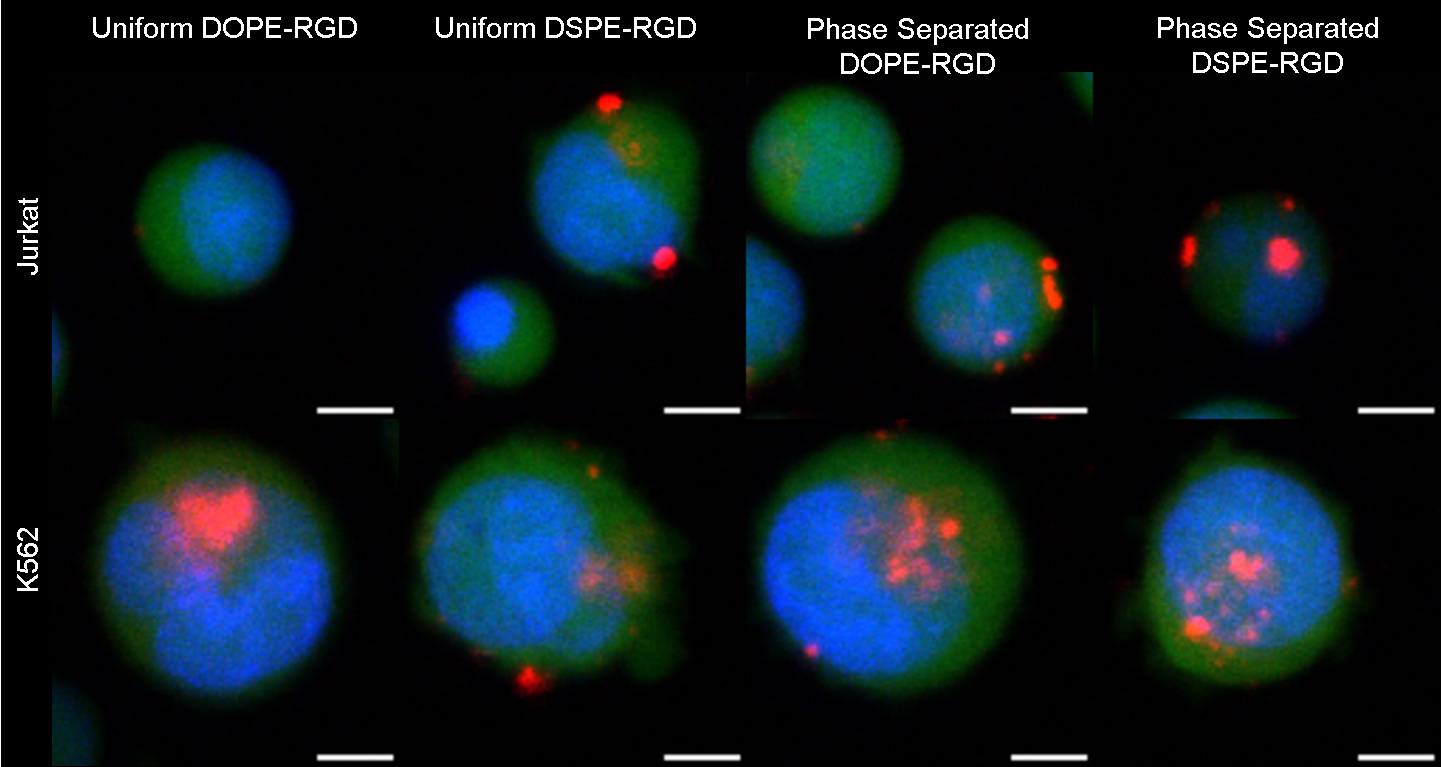

**Figure S7.** Enlarged microscopy images of vesicle uptake to individual Jurkat and K562 cells taken using confocal microscopy near the center of the cell. Scale bar = 5 μm.

**SUPPORTING TABLES**

**Table S1.** Vesicle compositions used in Figure 1. DLS size, PDI, and zeta measurements are displayed with SEM.

| **Name** | **Lipid Composition** | **Diameter (nm)** | **PDI** | **Zeta (mV)** |
| --- | --- | --- | --- | --- |
| DOPC No NTA | 69 mol% DOPC  0 mol% DSPC  30 mol% Chol  1 mol% DSPE-PEG2K  0.1 mol% 18:1 NBD PE  **OR** 18:1 Liss Rhod PE | 160 | 0.22 | -1.5 |
| DSPC No NTA | 0 mol% DOPC  69 mol% DSPC  30 mol% Chol  1 mol% DSPE-PEG2K  0.1 mol% 18:1 NBD PE  **OR** 18:1 Liss Rhod PE | 210 | 0.26 | -2.2 |
| 1:1 No NTA | 35 mol% DOPC  34 mol% DSPC  30 mol% Chol  1 mol% DSPE-PEG2K  0.1 mol% 18:1 NBD PE  **OR** 18:1 Liss Rhod PE | 170 | 0.17 | -1.5 |
| 2:1 No NTA | 23.3 mol% DOPC  45.7 mol% DSPC  30 mol% Chol  1 mol% DSPE-PEG2K  0.1 mol% 18:1 NBD PE  **OR** 18:1 Liss Rhod PE | 170 | 0.17 | -2.0 |
| 3:1 No NTA | 17.5 mol% DOPC  51.5 mol% DSPC  30 mol% Chol  1 mol% DSPE-PEG2K  0.1 mol% 18:1 NBD PE  **OR** 18:1 Liss Rhod PE | 190 | 0.25 | -2.2 |
| DOPC NTA | 69 mol% DOPC  0 mol% DSPC  30 mol% Chol  1 mol% DSPE-PEG2K  0.1 mol% 18:1 NBD PE  **OR** 18:1 Liss Rhod PE  +  1 mol% DGS-NTA(Ni) | 160 | 0.19 | -1.2 |
| DSPC NTA | 0 mol% DOPC  69 mol% DSPC  30 mol% Chol  1 mol% DSPE-PEG2K  0.1 mol% 18:1 NBD PE  **OR** 18:1 Liss Rhod PE  +  1 mol% DGS-NTA(Ni) | 190 | 0.24 | -2.5 |
| 1:1 NTA | 35 mol% DOPC  34 mol% DSPC  30 mol% Chol  1 mol% DSPE-PEG2K  0.1 mol% 18:1 NBD PE  **OR** 18:1 Liss Rhod PE  +  1 mol% DGS-NTA(Ni) | 170 | 0.21 | -2.2 |
| 2:1 NTA | 23.3 mol% DOPC  45.7 mol% DSPC  30 mol% Chol  1 mol% DSPE-PEG2K  0.1 mol% 18:1 NBD PE  **OR** 18:1 Liss Rhod PE  +  1 mol% DGS-NTA(Ni) | 170 | 0.18 | -3.0 |
| 3:1 NTA | 17.5 mol% DOPC  51.5 mol% DSPC  30 mol% Chol  1 mol% DSPE-PEG2K  0.1 mol% 18:1 NBD PE  **OR** 18:1 Liss Rhod PE  +  1 mol% DGS-NTA(Ni) | 180 | 0.18 | -2.1 |

**Table S2.** Vesicle compositions used in Figure 2. DLS size, PDI, and zeta measurements are displayed with SEM.

| **Name** | **Lipid Composition** | **Diameter (nm)** | **PDI** | **Zeta (mV)** |
| --- | --- | --- | --- | --- |
| Uniform 100:0 NTA | 68 mol% DOPC  0 mol% DSPC  30 mol% Chol  1 mol% DGS-NTA(Ni)  0.1 mol% 18:1 Liss Rhod PE  1 mol% DSPE-PEG2K  0 mol% DOPE-PEG2K | 160 | 0.20 | -2.0 |
| Uniform 75:25 NTA | 68 mol% DOPC  0 mol% DSPC  30 mol% Chol  1 mol% DGS-NTA(Ni)  0.1 mol% 18:1 Liss Rhod PE  0.75 mol% DSPE-PEG2K  0.25 mol% DOPE-PEG2K | 160 | 0.18 | -2.2 |
| Uniform 50:50 NTA | 68 mol% DOPC  0 mol% DSPC  30 mol% Chol  1 mol% DGS-NTA(Ni)  0.1 mol% 18:1 Liss Rhod PE  0.5 mol% DSPE-PEG2K  0.5 mol% DOPE-PEG2K | 160 | 0.19 | -1.3 |
| Uniform 25:75 NTA | 68 mol% DOPC  0 mol% DSPC  30 mol% Chol  1 mol% DGS-NTA(Ni)  0.1 mol% 18:1 Liss Rhod PE  0.25 mol% DSPE-PEG2K  0.75 mol% DOPE-PEG2K | 160 | 0.18 | -2.9 |
| Uniform 0:100 NTA | 68 mol% DOPC  0 mol% DSPC  30 mol% Chol  1 mol% DGS-NTA(Ni)  0.1 mol% 18:1 Liss Rhod PE  0 mol% DSPE-PEG2K  1 mol% DOPE-PEG2K | 160 | 0.13 | -2.4 |
| Phase-Separated 100:0 NTA | 17 mol% DOPC  51 mol% DSPC  30 mol% Chol  1 mol% DGS-NTA(Ni)  0.1 mol% 18:1 Liss Rhod PE  1 mol% DSPE-PEG2K  0 mol% DOPE-PEG2K | 170 | 0.12 | -2.2 |
| Phase-Separated 75:25 NTA | 17 mol% DOPC  51 mol% DSPC  30 mol% Chol  1 mol% DGS-NTA(Ni)  0.1 mol% 18:1 Liss Rhod PE  0.75 mol% DSPE-PEG2K  0.25 mol% DOPE-PEG2K | 190 | 0.17 | -2.6 |
| Phase-Separated 50:50 NTA | 17 mol% DOPC  51 mol% DSPC  30 mol% Chol  1 mol% DGS-NTA(Ni)  0.1 mol% 18:1 Liss Rhod PE  0.5 mol% DSPE-PEG2K  0.5 mol% DOPE-PEG2K | 180 | 0.11 | -2.0 |
| Phase-Separated 25:75 NTA | 17 mol% DOPC  51 mol% DSPC  30 mol% Chol  1 mol% DGS-NTA(Ni)  0.1 mol% 18:1 Liss Rhod PE  0.25 mol% DSPE-PEG2K  0.75 mol% DOPE-PEG2K | 200 | 0.20 | -2.0 |
| Phase-Separated 0:100 NTA | 17 mol% DOPC  51 mol% DSPC  30 mol% Chol  1 mol% DGS-NTA(Ni)  0.1 mol% 18:1 Liss Rhod PE  0 mol% DSPE-PEG2K  1 mol% DOPE-PEG2K | 200 | 0.22 | -1.6 |

**Table S3.** Vesicle compositions used for Figure 3.

| **Name** | **Lipid Composition** |
| --- | --- |
| Uniform 100:0 Biotin | 68 mol% DOPC  0 mol% DSPC  30 mol% Chol  1 mol% 18:1 Biotinyl Cap PE  0.1 mol% 18:1 Liss Rhod PE  1 mol% DSPE-PEG2K  0 mol% DOPE-PEG2K |
| Uniform 75:25 Biotin | 68 mol% DOPC  0 mol% DSPC  30 mol% Chol  1 mol% 18:1 Biotinyl Cap PE  0.1 mol% 18:1 Liss Rhod PE  0.75 mol% DSPE-PEG2K  0.25 mol% DOPE-PEG2K |
| Uniform 50:50 Biotin | 68 mol% DOPC  0 mol% DSPC  30 mol% Chol  1 mol% 18:1 Biotinyl Cap PE  0.1 mol% 18:1 Liss Rhod PE    0.5 mol% DSPE-PEG2K  0.5 mol% DOPE-PEG2K |
| Uniform 25:75 Biotin | 68 mol% DOPC  0 mol% DSPC  30 mol% Chol  1 mol% 18:1 Biotinyl Cap PE  0.1 mol% 18:1 Liss Rhod PE  0.25 mol% DSPE-PEG2K  0.75 mol% DOPE-PEG2K |
| Uniform 0:100 Biotin | 68 mol% DOPC  0 mol% DSPC  30 mol% Chol  1 mol% 18:1 Biotinyl Cap PE  0.1 mol% 18:1 Liss Rhod PE  0 mol% DSPE-PEG2K  1 mol% DOPE-PEG2K |
| Phase-Separated 100:0 Biotin | 17 mol% DOPC  51 mol% DSPC  30 mol% Chol  1 mol% 18:1 Biotinyl Cap PE  0.1 mol% 18:1 Liss Rhod PE  1 mol% DSPE-PEG2K  0 mol% DOPE-PEG2K |
| Phase-Separated 75:25 Biotin | 17 mol% DOPC  51 mol% DSPC  30 mol% Chol  1 mol% 18:1 Biotinyl Cap PE  0.1 mol% 18:1 Liss Rhod PE  0.75 mol% DSPE-PEG2K  0.25 mol% DOPE-PEG2K |
| Phase-Separated 50:50 Biotin | 17 mol% DOPC  51 mol% DSPC  30 mol% Chol  1 mol% 18:1 Biotinyl Cap PE  0.1 mol% 18:1 Liss Rhod PE  0.5 mol% DSPE-PEG2K  0.5 mol% DOPE-PEG2K |
| Phase-Separated 25:75 Biotin | 17 mol% DOPC  51 mol% DSPC  30 mol% Chol  1 mol% 18:1 Biotinyl Cap PE  0.1 mol% 18:1 Liss Rhod PE  0.25 mol% DSPE-PEG2K  0.75 mol% DOPE-PEG2K |
| Phase-Separated 0:100 Biotin | 17 mol% DOPC  51 mol% DSPC  30 mol% Chol  1 mol% 18:1 Biotinyl Cap PE  0.1 mol% 18:1 Liss Rhod PE  0 mol% DSPE-PEG2K  1 mol% DOPE-PEG2K |

**Figure S4.** Vesicle compositions used in Figure 4. DLS size, PDI, and zeta measurements are displayed with SEM.

| **Name** | **Lipid Composition** | **Diameter (nm)** | **PDI** | **Zeta (mV)** |
| --- | --- | --- | --- | --- |
| Uniform No RGD | 68.9 mol% DOPC  0 mol% DSPC  30 mol% Chol  0 mol% DSPE-RGD  0 mol% DOPE-PEG2K  1 mol% DSPE-PEG2K  0.1 mol% 18:1 Liss Rhod PE  0 mol% DOPE-RGD  0 mol% DSPE-RGD | 180 | 0.20 | -4.6 |
| Phase-Separated No DOPE-RGD | 23.2 mol% DOPC  45.7 mol% DSPC  30 mol% Chol  0 mol% DOPE-PEG2K  1 mol% DSPE-PEG2K  0.1 mol% 18:1 Liss Rhod PE  0 mol% DOPE-RGD  0 mol% DSPE-RGD | 160 | 0.19 | -8.7 |
| Phase-Separated No DSPE-RGD | 45.6 mol% DOPC  23.3 mol% DSPC  30 mol% Chol  1 mol% DOPE-PEG2K  0 mol% DSPE-PEG2K  0.1 mol% 18:1 Liss Rhod PE  0 mol% DOPE-RGD  0 mol% DSPE-RGD | 150 | 0.093 | -4.4 |
| Uniform DOPE-RGD | 58.9 mol% DOPC  0 mol% DSPC  30 mol% Chol  0 mol% DSPE-RGD  0 mol% DOPE-PEG2K  1 mol% DSPE-PEG2K  0.1 mol% 18:1 Liss Rhod PE  10 mol% DOPE-RGD  0 mol% DSPE-RGD | 190 | 0.17 | -4.7 |
| Uniform DSPE-RGD | 58.9 mol% DOPC  0 mol% DSPC  30 mol% Chol  0 mol% DSPE-RGD  1 mol% DOPE-PEG2K  0 mol% DSPE-PEG2K  0.1 mol% 18:1 Liss Rhod PE  0 mol% DOPE-RGD  10 mol% DSPE-RGD | 140 | 0.14 | -5.9 |
| Phase-Separated DOPE-RGD | 13.2 mol% DOPC  45.7 mol% DSPC  30 mol% Chol  0 mol% DOPE-PEG2K  1 mol% DSPE-PEG2K  0.1 mol% 18:1 Liss Rhod PE  10 mol% DOPE-RGD  0 mol% DSPE-RGD | 170 | 0.11 | -8.8 |
| Phase-Separated DSPE-RGD | 45.6 mol% DOPC  13.3 mol% DSPC  30 mol% Chol  1 mol% DOPE-PEG2K  0 mol% DSPE-PEG2K  0.1 mol% 18:1 Liss Rhod PE  0 mol% DOPE-RGD  10 mol% DSPE-RGD | 150 | 0.12 | -8.2 |

**Table S5.** DGS-NTA(Ni) vesicle compositions used for FRET assays in Figure S2.

| **Name** | **Lipid Composition** |
| --- | --- |
| Uniform 100:0 NTA FRET | 68 mol% DOPC  0 mol% DSPC  29 mol% Chol  1 mol% DGS-NTA(Ni)  0.5 mol% 18:1 Liss Rhod PE  0.5 mol% 16:0 NBD PE  1 mol% DSPE-PEG2K  0 mol% DOPE-PEG2K |
| Uniform 75:25 NTA FRET | 68 mol% DOPC  0 mol% DSPC  29 mol% Chol  1 mol% DGS-NTA(Ni)  0.5 mol% 18:1 Liss Rhod PE  0.5 mol% 16:0 NBD PE  0.75 mol% DSPE-PEG2K  0.25 mol% DOPE-PEG2K |
| Uniform 50:50 NTA FRET | 68 mol% DOPC  0 mol% DSPC  29 mol% Chol  1 mol% DGS-NTA(Ni)  0.5 mol% 18:1 Liss Rhod PE  0.5 mol% 16:0 NBD PE  0.5 mol% DSPE-PEG2K  0.5 mol% DOPE-PEG2K |
| Uniform 25:75 NTA FRET | 68 mol% DOPC  0 mol% DSPC  29 mol% Chol  1 mol% DGS-NTA(Ni)  0.5 mol% 18:1 Liss Rhod PE  0.5 mol% 16:0 NBD PE  0.25 mol% DSPE-PEG2K  0.75 mol% DOPE-PEG2K |
| Uniform 0:100 NTA FRET | 68 mol% DOPC  0 mol% DSPC  29 mol% Chol  1 mol% DGS-NTA(Ni)  0.5 mol% 18:1 Liss Rhod PE  0.5 mol% 16:0 NBD PE  0 mol% DSPE-PEG2K  1 mol% DOPE-PEG2K |
| Phase-Separated 100:0 NTA FRET | 17 mol% DOPC  51 mol% DSPC  29 mol% Chol  1 mol% DGS-NTA(Ni)  0.5 mol% 18:1 Liss Rhod PE  0.5 mol% 16:0 NBD PE  1 mol% DSPE-PEG2K  0 mol% DOPE-PEG2K |
| Phase-Separated 75:25 NTA FRET | 17 mol% DOPC  51 mol% DSPC  29 mol% Chol  1 mol% DGS-NTA(Ni)  0.5 mol% 18:1 Liss Rhod PE  0.5 mol% 16:0 NBD PE  0.75 mol% DSPE-PEG2K  0.25 mol% DOPE-PEG2K |
| Phase-Separated 50:50 NTA FRET | 17 mol% DOPC  51 mol% DSPC  29 mol% Chol  1 mol% DGS-NTA(Ni)  0.5 mol% 18:1 Liss Rhod PE  0.5 mol% 16:0 NBD PE  0.5 mol% DSPE-PEG2K  0.5 mol% DOPE-PEG2K |
| Phase-Separated 25:75 NTA FRET | 17 mol% DOPC  51 mol% DSPC  29 mol% Chol  1 mol% DGS-NTA(Ni)  0.5 mol% 18:1 Liss Rhod PE  0.5 mol% 16:0 NBD PE  0.25 mol% DSPE-PEG2K  0.75 mol% DOPE-PEG2K |
| Phase-Separated 0:100 NTA FRET | 17 mol% DOPC  51 mol% DSPC  29 mol% Chol  1 mol% DGS-NTA(Ni)  0.5 mol% 18:1 Liss Rhod PE  0.5 mol% 16:0 NBD PE  0 mol% DSPE-PEG2K  1 mol% DOPE-PEG2K |

**Table S6.** RGD vesicle compositions used for FRET assays in Figure S3.

| **Name** | **Lipid Composition** |
| --- | --- |
| Uniform DOPE-RGD | 58 mol% DOPC  0 mol% DSPC  30 mol% Chol  0 mol% DSPE-RGD  0 mol% DOPE-PEG2K  1 mol% DSPE-PEG2K  0.5 mol% 18:1 Liss Rhod PE  0.5 mol% 16:0 NBD PE  10 mol% DOPE-RGD  0 mol% DSPE-RGD |
| Uniform DSPE-RGD | 58 mol% DOPC  0 mol% DSPC  30 mol% Chol  0 mol% DSPE-RGD  1 mol% DOPE-PEG2K  0 mol% DSPE-PEG2K  0.5 mol% 18:1 Liss Rhod PE  0.5 mol% 16:0 NBD PE  0 mol% DOPE-RGD  10 mol% DSPE-RGD |
| Phase-Separated DOPE-RGD | 12.8 mol% DOPC  45.2 mol% DSPC  30 mol% Chol  0 mol% DOPE-PEG2K  1 mol% DSPE-PEG2K  0.5 mol% 18:1 Liss Rhod PE  0.5 mol% 16:0 NBD PE  10 mol% DOPE-RGD  0 mol% DSPE-RGD |
| Phase-Separated DSPE-RGD | 45.2 mol% DOPC  12.8 mol% DSPC  30 mol% Chol  1 mol% DOPE-PEG2K  0 mol% DSPE-PEG2K  0.5 mol% 18:1 Liss Rhod PE  0.5 mol% 16:0 NBD PE  0 mol% DOPE-RGD  10 mol% DSPE-RGD |
